# Investigating the Inhibitory Effects of Blebbistatin on Actomyosin Interactions in Myofibrils and Isolated Myofilaments

**DOI:** 10.1101/2025.02.17.638391

**Authors:** Daren Elkrief, Yu-Shu Cheng, Dilson E. Rassier

## Abstract

Myosin II is the molecular motor responsible for muscle contraction. Myosin II hydrolyses ATP into P_i_ and ADP, to convert chemical energy into mechanical work while attached to actin filaments. The relation between force generation and P_i_ release remains unclear. Many studies use chemical substances, such as blebbistatin, to study the transitions during actomyosin interactions. Blebbistatin and its derivatives selectively inhibit the actin-activated ATPase of myosin II, accumulating myosin cross-bridges in a pre-power-stroke state. Although the effects of blebbistatin have been explored, it is still unclear how blebbistatin affects force generation and the velocity of contraction. In this study, we used individual myofibrils, myosin and actin filaments, and isolated heavy meromyosin (HMM) and actin filaments to characterize the effects of blebbistatin. We observed that increasing concentrations of blebbistatin (i) decreased the force produced by myofibrils and isolated myosin filaments, (ii) decreased the maximum velocity of shortening produced by myofibrils and the myosin-induced actin sliding velocity, (iii) decreased the curvature of the force-velocity relation in a dose-dependent manner. Furthermore, UV radiation reduced the effect of blebbistatin, which was partially reversed if blebbistatin was bound to myosin before exposure to UV light. These results show that blebbistatin alters force and velocity generation at the molecular, myofilamentous and myofibrillar levels. This study has interesting implications in fields which rely on using blebbistatin to study cellular processes and confirms several results published in different experimental arrangements. Thus, this study is exploratory and confirmatory, and the findings have utility surrounding cell migration, muscle biophysics, cellular reproduction, or any processes that rely on the action of myosin II.

## Introduction

The molecular motor myosin II hydrolyses ATP into P_i_ and ADP, converting chemical energy into mechanical work while attached to actin filaments. This actomyosin interaction ultimately leads to force generation and muscle contraction. As such, altered myosin function contributes to many diseases (33, 61, 68) and weakness in aging (31), thus driving the development of myosin-targeted therapeutics (19, 33, 37, 61, 71). Unfortunately, the development of therapeutics is limited by the current understanding of the steps during actomyosin interactions, as some aspects of the cycle are still conflicting (5, 12, 13, 26, 34, 43, 44, 52, 60, 63, 72). Particularly, the relation between force generation and P_i_ release remains controversial and has spurred intense debate in the literature (12, 13, 26, 34, 43, 52, 60, 62, 72).

Several chemical substances have been used to investigate the transition state of the crossbridge cycle, resulting in changes to the ATPase, force generation and contractile velocity (1, 18, 28, 32, 38, 40, 51, 54, 55, 59, 63, 67, 69, 73). One advantage of using chemical substances instead of inducing mutations in myosin is that chemicals can be easily manipulated and used in different experimental preparations, from single molecules to contractile cells investigated *in situ*. A substance of value in this context is blebbistatin, whose binding site on myosin is well characterized (Fig. 1). This compound and its derivatives selectively inhibit the actin-activated ATPase of myosin II (28, 32, 51, 67). As a result, it has been suggested that blebbistatin causes an accumulation of cross-bridges in a pre-power-stroke state, with ADP and P_i_ at the active site (40, 42).

**Figure 1:**
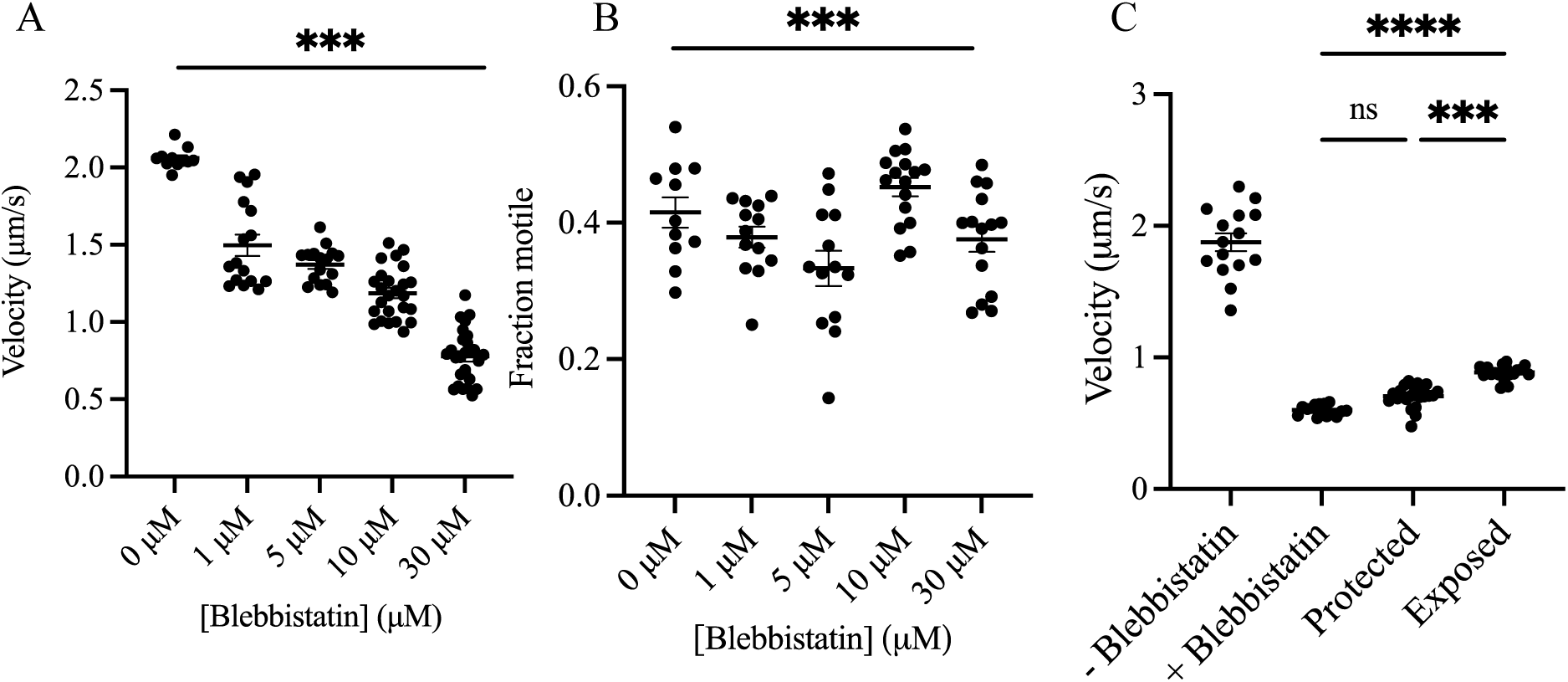
In vitro motility assay examining the effects of blebbistatin and UV light on filament velocity. A) Concentration-dependent decrease in filament velocity with increasing concentrations of blebbistatin. B) Variation in motile fraction across different blebbistatin concentrations. There were differences between motile fractions in each condition, although no pattern emerged (ANOVA P < 0.001). C) Effects of UV light on blebbistatin’s action. When 100 μM blebbistatin is first incubated with HMM for 30 minutes (protected) and then exposed to 488 nm light for 3 minutes, the decrease in average velocity is insignificant compared to using 100 μM blebbistatin in the dark, indicating significant blebbistatin protection (p = 0.08). However, filaments with 100 μM blebbistatin first UV-treated and then incubated with HMM (exposed) show a significantly higher velocity than those incubated with 100 μM blebbistatin in the dark (p < 0.001). Conditions from left to right are as follows: 1) No blebbistatin, 2) Blebbistatin, 3) Blebbistatin + HMM, then UV (protected), 4) UV + Blebbistatin, then HMM (exposed). Data are presented as mean ± S.E.M.

The mechanism by which blebbistatin inhibits contractility is not simple, as it specifically targets the step in the actomyosin cycle that is poorly understood, i.e., the P_i_ release and force generation. There are also results that are difficult to reconcile; blebbistatin decreases the isometric force in muscles fibers independent of myosin regulatory light chains (RLCs) phosphorylation (64), accompanied by a smaller decrease in stiffness (42, 49), but it decreases the unloaded shortening velocity only when RLCs are phosphorylated (64). Blebbistatin stabilizes the start of the power-stroke even without P_i_ at the active site (69), which makes interpretation of its effects challenging. Therefore, there is the question: how does blebbistatin affect the force and velocity of contraction at the filamentous and molecular levels? Importantly, we further this query by asking whether the effects of blebbistatin translate across *in vitro* motility and myofilament experimental procedures.

In this study, we used three preparations to characterize the effects of blebbistatin in skeletal muscle myosin: individual myofibrils, isolated myosin-actin filaments, and isolated heavy meromyosin (HMM)-actin filaments. Myofibrils represent the smallest muscle structure that maintains the three-dimensional lattice intact, with the major contractile and regulatory proteins. They allow rapid activating and relaxing of the contractile system, and direct measurement of sarcomere length and specific force, given its small cross-sectional area (∼1 μm). They also allow the use of rapid length changes for measurements of the force-velocity relation (21, 45, 57, 58). The myosin-actin filaments system, on the other hand, allows us to directly evaluate the effects of blebbistatin on the force produced at the molecular level during actomyosin interactions (6, 7, 15, 25). Isolated HMM allows measurements of the myosin-driven speed of actin motility of the thin filament (6, 7, 15, 25).

## Methods

### Animals

Experiments were performed with isolated myofibrils from the rabbit psoas muscle, native thick filaments isolated from the rabbit psoas muscle, and native thick filaments isolated from the anterior byssus retractor muscle (ABRM) of *Mytilus edulis* (mussel). The ethical protocol for use of animal material was approved by the Animal Care Committee at McGill University (Ref. No. MCGL-5227) and the Canadian Council on Animal Care.

### Experiments with isolated myosin molecules and filaments

#### Myosin and HMM preparation for the in vitro motility assay (IVMA)

For measurements using the IVMA, skeletal muscle myosin was isolated from rabbit psoas samples as previously done in our laboratory (6, 7, 15, 24, 25). Briefly, 5 g of muscle was homogenized in 15 mL of Hasselbalch-Schneider buffer A (0.1M KH2PO4/K2HPO4 at pH 6.4; 0.6M KCl, 10mM Na_4_P_2_O_7_·10H_2_O, 1mM MgCl_2_, and 20mM ethylene glycol tetra-acetic acid) using an Omni mixer homogenizer (Omni International, Inc., GA). The solution was stirred at 4°C for 15 min and the reaction was stopped by adding 20 mL of distilled water at 4°C. The mixture was centrifuged at 3,000 g (5804 R, Eppendorf AG, Hamburg, Germany) for 10 min, and the myosin supernatant was passed through a 55-mm hardened circle filter paper (No. 54, GE Healthcare, IL). The filtrate was diluted with 4°C distilled water and centrifuged at 10,000g for 15 min. The pellet was washed with buffer B (20mM K_2_HPO_4_ at pH 7.2; 0.12M KCl; 1mM EDTA; and 1mM dl-dithiothreitol (DTT)), resuspended in buffer C (50mM sodium pyrophosphate and 1mM DTT at pH 7.8), and centrifuged at 10,000 g for 10 min. The supernatant was filtered through No.54 filter paper (Whatman, GE), and the myosin solution was stored in glycerol in a −80°C freezer, or else used for HMM preparation.

An imidazole-based method for HMM preparation was adapted from previous studies (6, 7, 15, 24, 25). Nine volumes of buffer solution (0.1 M imidazole hydrochloride, 0.1 mM EGTA, and 2 mM DTT added after setting to pH 7.0) were added to a centrifuge tube containing 21 mg of stock myosin solution, mixed slowly to precipitate myosin filaments, and then left on ice for 10 min. The solution was centrifuged at 15,000 g for 60 min at 4°C. The pellet was dissolved in buffer (20 mM imidazole hydrochloride, 1 M KCl, 4 mM MgCl_2_, and 10 mM DTT added as before) 2 times to obtain a final concentration of 15 mg/mL myosin in buffer. The myosin solution was then incubated in a shaking water bath at 25°C for 5 min. 1 mg/mL chymotrypsin was added to the myosin to a final concentration of 12.5 μg/mL, and this mixed solution was set in a water bath for 10 min at 25°C. Seven microliters of 0.2 M stock PMSF protease inhibitor and 12.6 mL of cold buffer were added to the solution and gently mixed. The mixture was left on ice for up to 1 h and then centrifuged at 15,000 g for 60 min at 4°C. At this stage, the myosin was pelleted, and HMM was recovered from the supernatant. The HMM concentration was determined via the Bradford assay (Bio-Rad, CA). Sucrose was added to aliquots of HMM in Eppendorf tubes to a final concentration of 2 mg/mL and stored at −80°C in liquid nitrogen.

#### Myosin filaments isolation for force measurements

For force measurements, fresh thick myosin filaments were obtained from the anterior byssus reactor muscle of *Mytilus edulis* (mussel) using a previously described protocol (6, 7, 15, 24, 25). Mussels (Mytilus edulis) were purchased from a local seafood market and brought to the laboratory on the day of filament extraction. Pairs of byssus reactor muscles were quickly excised from fresh mussels and immediately placed in thick filament buffer solution [10 mM PIPES (pH 7.0), 10 mM MgCl_2_, 2 mM EGTA, 10 mM ATP, 2 mM DTT]. The filaments were dissected by using a scalpel (No. 10 surgical blade equipped; FEATHER) and washed with rigor solution [50 mM Tris, 100 mM KCl, 2 mM MgCl_2_, and 1 mM EGTA (pH 7.0)]. The filaments were teased apart and isolated by using surgical tweezers (No. 5 fine-needle sharp; Sigma-Aldrich), and anterior byssus retractor muscles (ABRMs) were isolated from fresh Mytilus edulis by cutting with micro-dissecting scissors (size 4, curved, sharp point stainless steel). After extraction, the samples were stored on ice at 4°C in rigor solution. After 24 h, these muscles were transferred to a mixed solution containing rigor/glycerol (50%/50%) and stored at −20°C. Prior to experimentation, muscle bundle strips were then homogenized (SNMX 1092, Omni Inc., GA) three times for 7s, with 1-minute intervals, and then placed on ice. The muscle homogenate was mixed with an equal volume of the thick filament buffer solution containing 0.1% Triton X-100 and left on ice for 15 minutes. The homogenate was precipitated in a centrifuge at 700 g for 5 min, and the pellet was discarded. The supernatant was further centrifuged at 4,500 g for 40 minutes, and pelleted filaments were resuspended in the thick filament buffer. Centrifugation steps were repeated, and thick filaments were resuspended in an ATP-free buffer solution.

#### F-actin preparation for the IVMA and the force measurement experiments

Actin was extracted from rabbit psoas and isolated according to a modified protocol used previously in our laboratory (6, 7, 15, 24, 25). Briefly, actin was purified in 10 mL of fresh G-actin buffer (2 mM imidazole, 0.2 mM Na_2_ATP, and 0.2 mM CaCl_2_) per gram of acetone powder. The solution was stirred for 30 minutes on ice and filtered through four layers of cheesecloth. The liquid was further extracted with 10 mL of G-actin buffer per gram of acetone powder and stirred for 10 minutes at 0–0.5°C. Filtrates were centrifuged at 20,400 *g* for 20 minutes at 4°C and passed through a cheesecloth, and pellets were discarded. The supernatant was then dissolved in 50 mM MgCl_2_, and actin filaments were polymerized with the addition of 0.2 mM ATP. The solution was left for 1 h at room temperature while KCl was slowly added to a final concentration of 0.8 M. The solution was stirred again for 10 minutes and then centrifuged at 12,000 *g* for 1 hr, at 4°C, after which the supernatant was discarded. Actin concentration of 7 μM was determined using absorbance measurement at 290 nm using E = 0.63 mL/mg × cm, as described elsewhere. The product purity was determined using 10% SDS-PAGE. HMM and polymerized actin filaments (F-actin) were frozen in liquid nitrogen in the presence of 2 mg/mL sucrose and stored at −80°C, ideal conditions for long-time storage.

#### Blebbistatin

Blebbistatin (+/-) (Sigma-Aldrich) was dissolved stored in dimethylforamide (DMF) to 20 mM and stored at –20°C before use. On the day of the experiments, blebbistatin was dissolved to the desired concentrations listed in each experiment in their respective sections. Concentrations ranging from 1 to 30 μM were effective at either slowing motility or reducing force with a similar half maximum effectiveness expected from a racemic blebbistatin (41, 50). Unless otherwise noted, blebbistatin was shielded from all light sources and experiments were performed quickly to avoid blebbistatin degradation.

#### Force measurements and micro fabricated cantilevers

Force measurements in myofilaments were conducted in an experimental chamber mounted on the stage of an inverted microscope (Nikon TE2000-U) equipped for dark-field illumination, bright-field illumination, and fluorescence imaging. Dark-field illumination was used to image myosin filaments (22, 23). Fluorescence microscopy was used to image actin filaments with a filter set for Alexa-488 (Exciter ET470/40x, Dichroic T495LP, Emitter ET525/50m; Chroma). Images were captured with a Rolera-MGi Plus video camera (QImaging) and recorded using Streampix4 software (pixel size: 120 nm; collection rate: 50 fps; Norpix). The combination of dark-field microscopy and fluorescence microscopy allowed the simultaneous imaging of both myosin and actin filaments.

Microfabricated cantilevers were used for force measurements during myosin-actin interactions (3, 22, 23). The cantilevers were made from 400 μm-thick silicon nitride wafer, followed by photolithography process, as previously used in our laboratory (6, 15). The cantilever tips were coated with a 50 nm platinum coating on the tip to improve reflectivity and optical contrast. The dimensions of flexible cantilevers were chosen to provide stiffness in the ranges of forces to be measured during the experiments (length: 550 μm; width: 1 μm; thickness: 0.6 μm). The stiffness of the cantilevers was obtained using a resonance frequency detection method, as previously described (10). Briefly, the cantilever tip was illuminated with a laser, and the absorption peak was detected on a single photodiode detector (PIN-5D; UDT Sensors, Inc., Hawthorne, CA).

Mechanical vibration was induced on the cantilever with a piezoelectric motor. When the vibration of the cantilever reached the maximum amplitude response, the resonance frequency was measured. With the known resonance frequency, the elasticity modulus and the stiffness of the cantilevers were calculated. The stiffness obtained with this calibration method is not specific to a particular point on the cantilever tip but the average stiffness for the cantilever tip. The stiffness of the cantilever is 0.18 pN/nm in this study.

The cantilevers were glued on the bottom of the metal holders, which were connected to micromanipulators that allow three-dimensional manipulation inside of the experimental chamber (Fig. 1C). Although native thick and actin filaments have a small degree of compliance (30, 36), filament stretching or shortening was not observed in these experiments (Fig. 1D) within the resolution of our system (120-nm pixel size when the overlap was measured). Consequently, the degree of filament overlap should be proportional to the cantilever displacement. The cantilevers were placed in the experimental chamber separated ∼10–15 mm apart. 10 μL of myosin filament solution was added near the rigid cantilever, and 10 μL of α-actinin solution was added near the flexible cantilever, followed by a 10-minute incubation period. A flow of standard AB/BSA/GOC/ATP solution (AB: 0.5 mg/mL BSA, 0.018 mg/mL catalase, 0.1 mg/mL glucose oxidase, 3 mg/mL glucose, 20 mM DTT, and ATP concentrations between 50 and 2,000 μM) was injected into the chamber with a syringe pump (Pump 33; Harvard Apparatus) at a speed of 0.5 mL/minute. This flow washed the excess of α-actinin and myosin filaments. After 2 minutes, the flexible and the rigid cantilever were moved into the field of view (×100 magnification). Fluorescently labeled actin filaments (concentration: 2–4 nM) were injected into the chamber, and the flow facilitated their spontaneous attachment to one of the α-actinin-coated cantilevers. Blebbistatin was allowed to interact with thick filaments in a dark-enclosed test tube for 30 minutes prior to usage. The number of measurements taken (N = 10 controls, N = 23 blebbistatin in total, N = 11 Blebbistatin + UV treatment) are represented by the number of actin-myosin cross bridges which were successfully formed. While may more cross bridges may have been formed than those reported in this study, a subset of these were recorded due to the technical difficulty in capturing the correct cross bridge. For the experiments using blebbistatin, blebbistatin was added to AB/BSA/GOC/ATP solution at 0, 1, 5, 7.5, 10, or 12.5 μM concentrations. Data collection was done as quickly as possible to avoid unwanted UV damage. The flow was maintained constantly to align the actin filaments ∼90° perpendicular to the cantilevers. Myosin filaments were then injected into the chamber and adhered spontaneously to the cantilevers. All the experiments were performed at room temperature.

One actin filament and one myosin filament attached at the cantilever’s tips, at a distance <50 μm from the tip, were chosen for mechanical experimentation. Using the micromanipulators, the cantilevers were brought to proximity until they interacted (Supplementary Video 1). Once the filaments interacted, they initiated force production and consequently displacement of one of the cantilevers, which was tracked with Image software (National Institutes of Health). The force (F) during interactions was calculated from the displacements of the cantilevers, as explained previously (3, 22, 23): F = k × Δd, where k is stiffness and Δd is the amount of cantilever displacement. At times, the alignment of the filaments was not at a 90° angle to the cantilever; in these cases, the force component on the vertical axis during cantilever displacement was enhanced by an angular component represented by the vertical axis. The full force generated by the filaments during interaction was then adjusted such that F = Fx + Fy, where Fx is the vector component of force along the horizontal axis and Fy the vector component of force along the vertical axis. The images of the cantilever displacements were analyzed using an automatic algorithm (ABSnake for ImageJ; National Institutes of Health). The system with cantilevers can be servo-controlled either in position or to be used for force feedback. We were able to induce changes of ∼200 nm without significantly increasing the noise to the system.

#### In Vitro Motility Assay

Motility experiments were conducted as previously performed in our laboratory (6, 7, 15, 24, 25). Glass coverslips (60 × 24 mm^2^, No. 0, Menzel-Glaser, Braunschweig, Germany) were soaked in 70% EtOH overnight, dried, and coated with 1% nitrocellulose to create an isotropic and nonspecific binding surface for HMM attachment. Flow cells were constructed using double-sided adhesive tape to build a fluid chamber between a small coverslip (20 × 20 mm^2^) and a large coverslip (60 × 24 mm^2^). One different coverslip was used for each condition, and samples were taken at least five times in different areas of the coverslip.

The assay was developed by slowly adding 60 μL volumes of the following solutions to the side of the flow chamber in the following order: (1) HMM (120 μg/mL) diluted to 60 μg/mL in L65 solution (low ionic strength solution (LISS), composed of 1 mM MgCl_2_, 10 mM MOPS, 0.1 mM K_2_EGTA, pH 7.4, 1 M KCl, and 15 mL DTT), (2) bovine serum albumin (BSA, 1 mg/mL) in LISS, (3) L65, (4) fluorescently labeled actin diluted to 10 nM in L65, (5) L65, and (6) assay solution (15 mM LISS, 1 mM MgATP, 10 mM DTT, 0.64% methylcellulose, 130 mM KCl, 3 mg/mL glucose, 0.1 mg/mL glucose oxidase, 0.02 mg/mL catalase, 2.5 mM phosphocreatine, and 0.2 mg/mL creatine kinase). At each step, blebbistatin was added from a stock solution of 20 mM to final concentrations of 0, 1, 5, 10 and 30 μM in the HMM, BSA, L65 and assay solutions for 30 minutes prior to usage in the IVMA. Unless otherwise stated, all IVMA preparation steps were performed in the dark to minimize blebbistatin damage. To maximize equilibration, solutions at the following steps were allowed to sit for longer on the coverslip: HMM (5 minutes) BSA (2-5 minutes), L65 (15 minutes). For UV experiments, HMM in blebbistatin tubes were split into two tubes, one dark and one exposed to light. The light tube was incubated for 3 minutes under the UV light. Light exposed blebbistatin was then used for light/dark comparison studies at the desired concentrations.

The sliding actin filaments were “mage’ under an inverted darkfield fluorescent microscope (Axio Observer D1, Zeiss, Jena, Germany, or Eclipse TE300, Nikon, Tokyo, Japan) using a ×100 objective. Images were 1,200 × 1,200 pixels and 110 nm/pixel for the ×100 objective (frame rate: 10 fps). MatLab (MatLab R2017b; MathWorks, Natick, MA) software was used to obtain the velocity of filament sliding. Videos were acquired as quickly as possible, within 5 minutes of placing assay solution on the coverslip, to avoid increasing variance in motility over time. Immobile actin filaments as well as those with aberrant start and stop motion were excluded from analysis based on criteria as outlined elsewhere (15). Each data point represents the average of at least 15 filaments from 15 videos, pooled from three separate experiments. The average temperature was 25°C, and analysis was cut off at 0.2 μm/s to exclude non-motile and wiggling filaments.

### Experiments with myofibrils

Female New Zealand White Rabbits were obtained by Charles River Canada (QC, Canada) and were euthanized at an age between 4-12 months. They were fed ad libitum and weighed between 2.5 and 3.5 kg. They were euthanized with exsanguination under general anesthesia. Accordingly, a subcutaneous injection was administered (premedication in 0.1 mg/kg Glycopyrrolate, 0.5 mg/kg Butorphanol, and 0.75 mg/kg Acepromazine) followed by an intramuscular injection (anesthetics in 35 mg/kg Ketamine, 5 mg/kg Xylazine). An incision of ∼20 cm was made in the mid-section of the animals’ belly using a sharp scalpel (No. 10 surgical blade equipped: FEATHER). Strips of the psoas muscle (∼3 mm in diameter and ∼5 cm in length) were dissected using surgical tweezers (No. 5 fine-needle sharp; Sigma-Aldrich). The muscle strips were tied in 6″ applicator sticks (Fisher) with black braided silk (2-0, SP118, LOOK) and stored on ice at −4°C in rigor solution. Approximately 6 h after extraction, the muscle samples were transferred to a mixed solution containing rigor/glycerol (50%/50%) and stored at −20°C.

#### Sample preparation

Myofibrils were obtained following procedures previously used in our laboratory (8, 14, 20, 21, 39, 57, 58). On the day of the experiment, the sample was defrosted at 4°C in rigor solution for 30 min. Then, the samples were left in a relaxing solution [100 mM KCl, 10 mM PIPES (pH 7.0), 10 mM MgCl_2_, 2 mM EGTA, 10 mM ATP, and 2 mM DTT] for 1 h, before being diced into thin strips and homogenized on ice (SNMX 1092, OmniInc) at 2°C for 30 s, with 1-min intervals. The homogenate was centrifuged (5804 R, Eppendorf) at 4,500 *g* for 30 min to remove damaged myofibrils and other contaminants.

#### Myofibril solutions

The rigor solution (pH 7.0) used to store the myofibrils was composed of (in mM) 50 Tris, 100 NaCl, 2 KCl, 2 MgCl_2_, and 10 EGTA. The experimental solutions used during the experiments (pH 7.0) was composed of (in mM) 20 imidazole, 14.5 creatine phosphate, 7 EGTA, 4 MgATP, 1 free Mg^2+^, free Ca^2+^ ranging from 1 nM (pCa 9.0) to 32 μM (pCa 4.5), and KCl to adjust the ionic strength to 180 mM. The final concentrations of each metal-ligand complex were calculated using a computer program based on previous studies (13, 37).

#### Blebbistatin treatment

Blebbistatin was dissolved in dimethylformamide (DMF) to reach a concentration of 20 mM and was stored at −20°C before use. On the day of the experiment, 1 μL of blebbistatin was diluted in 4 mL of activating (pCa 4.5) or relaxing (pCa 9.0) solution to reach final concentrations of 2 μM and 5 μM. This concentration range is commonly used and allows for comparisons with other studies (9, 16, 42, 64). Furthermore, in previous experiments in our laboratory, we noted that 5 μM blebbistatin consistently decreased the force to levels between 40 % and 50 % of maximal force (42).

#### Instrumentation

The myofibrils were tested with a custom Atomic Force Microscope (AFM) as previously described (30, 53, 57, 58). Myofibrils were placed into rigor solution inside a chamber that was maintained at room temperature. An inverted microscope with phase-contrast illumination was used to measure the striation pattern on the myofibrils, based on the dark bands of myosin (A-bands) and the light bands of actin (I-bands). The pattern was used to measure their initial sarcomere length (SL). A myofibril was glued and suspended between the tips of a rigid glass microneedle and a silicon atomic force cantilever (ATEC-CONTPt-20; Nanosensors, Watsonville, CA). The microneedle was used to shorten the myofibril using a piezo-electric controller during experiments designed to measure the force-velocity relation. The cantilever stiffness was calculated following procedures previously used in our laboratory (57, 58), with a value of 42.2 nN/μm. Throughout the experiments, the surface of the experimental chamber was visualized by an inverted phase-contrast microscope (Eclipse TE2000-U) under high magnification (Nikon Plan Fluor, X100, numerical aperture 1.30 + 1.5 × microscope magnification). All experiments were video-recorded (Hamamatsu Orca-ER digital camera) for subsequent data analysis.

#### Experimental protocol

Contractions of myofibrils were induced by a fast-switching system that allowed exchanges between relaxing and activating solutions delivered from a double-barrelled perfusion pipette, that was directed at the selected myofibril, as described in detail elsewhere (20, 21, 53). The experiment started with the myofibrils being surrounded by a relaxing solution (pCa 9.0). The myofibrils were activated by quickly switching the position of the double-barreled pipette, which immersed the myofibril in an activating solution (pCa 4.5). When surrounded by a solution containing high Ca^2+^ concentration, the myofibrils produced force, causing deflection of the AFC. The deflection of the cantilever was recorded using an optical system developed in our laboratory (30). The force (F) was extracted from the cantilever displacement and calculated as *F* = *k**·**Δd*, where *k* is the cantilever stiffness and *d* is the cantilever displacement. Forces were normalized by the myofibril cross sectional area, assuming circular geometry.

Myofibrils were also tested for the maximal velocity of shortening, and for constructing a force-velocity relation for analysis of power output. First, the myofibrils underwent a slack test: they were allowed to reach maximal force, and a predetermined length step was imposed rapidly (2ms) allowing the myofibril to become slack and tension drop to zero. Force redeveloped over time proportionally to the length change. Four separate slack tests were performed on each myofibril corresponding to maximum sarcomere length (SL) steps of 15%SL, 20%SL, 25%SL and 30%SL. It should be noted that only isometric force was measured and not force transients or force kinetics due to experimental design constraints, technical limitations, and the focus on our main outcomes. Future studies could investigate activation and relaxation kinetics to provide a more comprehensive understanding of transient force responses in myofibrils. The data was plotted as the time required to re-develop tension relative to the imposed length step and was fitted with a first order least squares regression line. The corresponding slope of this fitted line represents the unloaded shortening velocity. Finally, the myofibrils were released during maximal activation by a step reduction in length (0.010SL/sec – 4 SL/sec). The force at the end of shortening was measured, and Hill’s equation was used to fit the force/velocity data for shortening for each myofibril: (F + a) × (V + b) = b (Fo + a), where a and b are constants, F is force at a given velocity, V is the velocity, and Fo is the maximal isometric force when V = 0. Vmax occurs when F = 0, such that Vmax = (b × Fo)/a).

### Statistical analysis

Data are presented as mean ± 95 % confidence limits unless otherwise stated. The force values of the experiments conducted with myofibrils were compared using a one-way analysis of variance (ANOVA) for repeated measures. The force, velocity of actin motility and fraction motility collected during the experiments with isolated filaments were also compared with one-way ANOVA for repeated measures. All statistical analyzes and curve fittings were performed using the GraphPad Prism software (Version 6.07; GraphPad software Inc, USA). A significance value of p < 0.05 was used for all analyses.

## Results

### In vitro motility assay

The IVMA was used to determine whether blebbistatin would affect the velocities of myosin-propelled actin filaments. In this study, we used a motility analysis algorithm where multiple filaments are averaged over the length of a video capture, over a minimum of 100 frames with 0.2 seconds per frame. Thus, our results show multiple filaments that are averaged over time, and the *N* represents the number of videos that we used to evaluate the velocity of actin motility.

The blebbistatin concentration used in these experiments ranged from 0 – 30 *μ*M. We observed a decrease in the actin sliding velocity with blebbistatin, in a dose dependent manner (Fig. 1). When we used 30 μM of blebbistatin, the actin sliding velocity dropped by ∼60%. The range at which blebbistatin reduced motility by 50% (IC_50_) in these experiments was in concentrations between 0.5-5 *μ*M, in line with previous results (32, 49). The motile fraction did not change significantly with increasing blebbistatin (Fig. 2B), showing it did not affect the number of filaments gliding over myosin.

**Figure 2.**
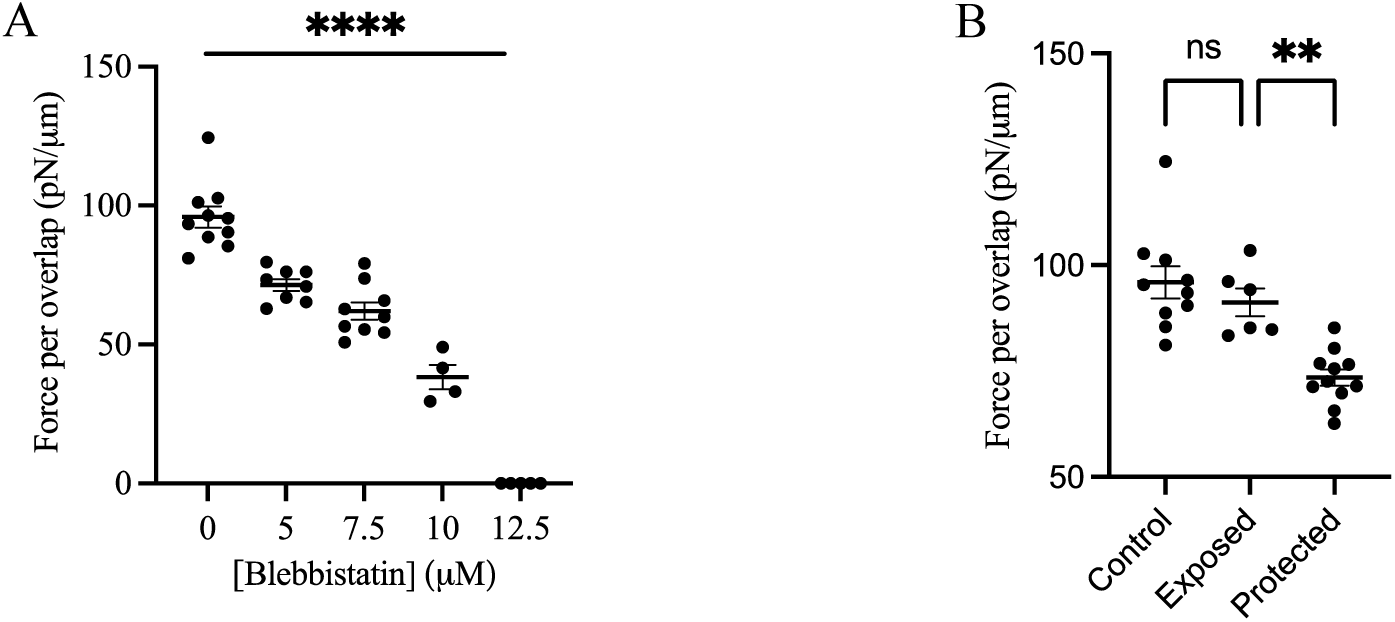
Filament force measurement. Actin and myosin filaments were made to interact. Force were recorded by cantilevers in a force measurement system. **A)** Forces normalized to overlap length between actin and myosin filaments were assessed with 0, 5, 7.5, 10, and 12.5 μM blebbistatin. Force decreased with increasing concentrations of blebbistatin, until no force was detected at 12.5 μM blebbistatin (ANOVA P < 0.0001). **B)** Filament force of blebbistatin administration subject to UV treatment before incubation with myosin (Exposed) and UV treatment after myosin incubation (Protected) at 10 μM blebbistatin. Force decreased only by ∼5% when blebbistatin was first incubated UV light (95% of the force relative to controls was maintained,) and then made to interact with myosin (p = 0.59). Force decreased by ∼23% after blebbistatin was protected compared to when it was exposed (Tukey’s multiple comparison, p = 0.0025). All values shown as mean ± S.E.M.

As with the force measurements, we hypothesized that UV would affect the blebbistatin-induced reduction on force. Indeed, there was a 52% decrease (1.88 μm/s ± 0.068, N = 15, control) in the acting sliding velocity when 100 μM blebbistatin was exposed to light before addition of HMM (0.89 ± 0.012 μm/s, N = 19 UV then incubate with HMM) (p < 0.001). When 100 μM blebbistatin was first incubated with HMM for 30 minutes and then exposed to 488 nm light for 180 seconds (0.707 ± 0.017 μm/s, N = 23, incubate with HMM then UV), there was an insignificant decrease in average velocity when compared to using 100 μM blebbistatin in the dark (0.60 ± 0.01 μm/s, N = 15, 100 μM) (p = 0.08), suggesting blebbistatin protection from UV light. There was also a higher velocity in filaments treated with 100 μM blebbistatin that were first UV-treated and then incubated with HMM compared to those which were incubated with 100 μM blebbistatin in the dark (p < 0.001). Interestingly, blebbistatin exposed to UV for 3 minutes showed the same reduction in efficacy to that exposed to incandescent laboratory light for 30 minutes, possibly indicating that there was an excess of blebbistatin:HMM (at 100 μM), and thus the maximum decrease in efficacy was reached during our experiments (Supplementary Fig. 1).

### Myofilament experiments

We used a filament force measurement system developed in our laboratory (6, 10, 24, 25) to determine the extent to which blebbistatin reduces forces produced by isolated myosin-actin filaments (Fig 2). We used the muscle myosin of the molluscan thick filaments (ABR) because of their length that is longer than skeletal filaments, making the experiments more feasible due to a larger overlap with actin during interaction (25). The force per overlap produced by thick filaments was 95.93 ± 3.81 pN/μm before treatment with blebbistatin (N = 10). These results corroborate results from previous studies that measured the force in isolated filaments (6, 10, 24, 25).

Blebbistatin decreased the forces developed by the filaments, that were ∼60% lower than control filaments at a concentration of 10 μM blebbistatin (38.29 ± 4.39, pN/μm N = 4), ∼35% lower at 7.5 μM blebbistatin (62.06 ± 3.15 pN/μm, N = 9), and ∼25% lower at 5 μM blebbistatin (71.45 ± 2.10 pN/μm, N = 8) (Fig. 2A)(ANOVA p < 0.001). There was no force produced (0 pN/μm) when the filaments were treated with 12.5 μM blebbistatin, although binding could still occur.

It is known that UV light reduces the effects of blebbistatin on myosin molecules (56). We used this property to test potential mechanisms of blebbistatin, following a rationale that HMM might protect blebbistatin from UV damage if the compound was inside the HMM pocket. We therefore assessed whether there was a difference between UV treatment when blebbistatin was in the HMM pocket versus UV treatment of blebbistatin out of the pocket, i.e., before blebbistatin was added to myosin. We hypothesized that UV effects on blebbistatin-induced reduction on force may be attenuated by the close interaction with the hydrophobic pocket of myosin. In fact, there was a decrease of 23.4% ± 2.01 in blebbistatin (10 μM) effectiveness when it was subjected to UV light for 180 seconds before incubation with myosin (73.45 ± 1.93 pN/μm, N = 11) compared to controls (Fig. 2 B, C). However, when UV treatment of blebbistatin was used after it had been incubated for 30 minutes with HMM, we observed a small decrease of 4.93% ± 3.42 in force (91.2 ± 3.28 pN/μm, N = 6) relative to controls, suggesting a significant protection (p < 0.001).

### Myofibril experiments

Myofibrils were experimented before and after blebbistatin treatment to test its effects on contractile properties. When activated the myofibrils at an average sarcomere length of 2.6 µm and at a temperature of 15°C, the force (normalized to area) produced by the myofibrils before blebbistatin treatment was 136.4 ± 15.2 nN/µm^2^ (Fig 3), a value that is close to values reported in previous studies in our laboratory (3, 9, 11, 14, 48, 53) and others (29, 47, 73, 75). When the myofibrils were treated with 2 µM and 5 µM of blebbistatin, the force per area decreased significantly to 64.5 ± 8.5 nN/µm^2^ and 48 ± 6.0 nN/µm^2^, a percentage decrease in the same range as published previously in other studies that used similar experimental conditions (9, 16, 42, 64), a result that was confirmed statistically (P = 0.01).

**Figure 3.**
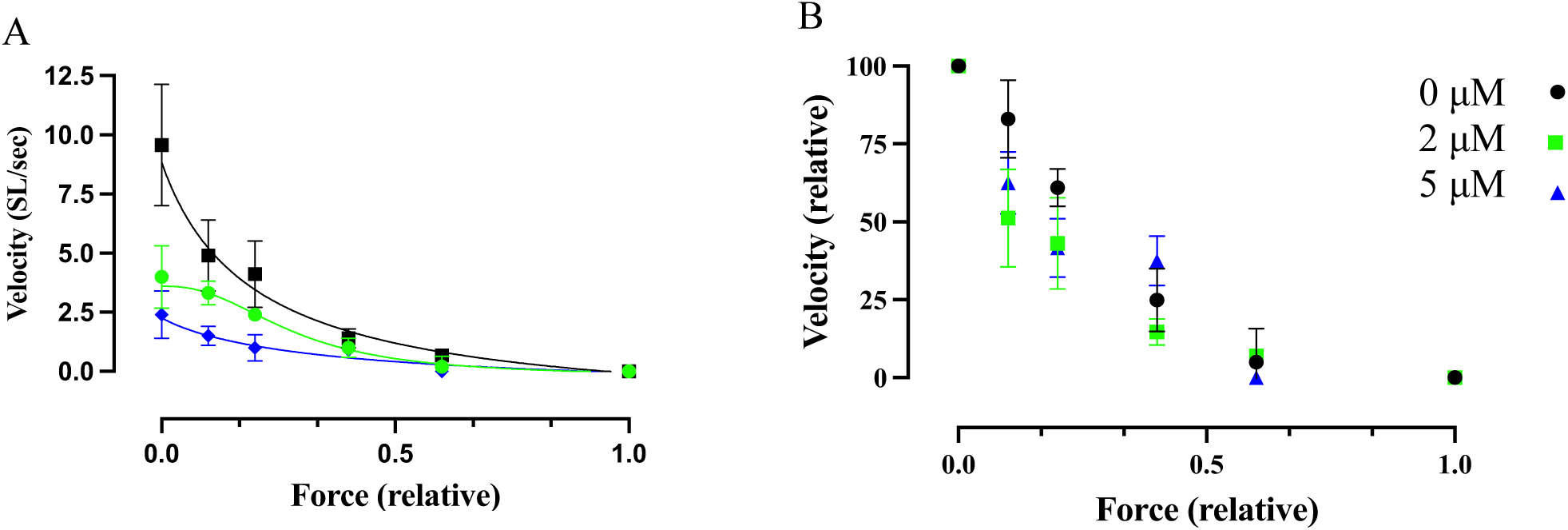
Force-velocity curves of myofibrils treated with 0 μM (black), 2 μM (green), and 5 μM (blue) blebbistatin. A) Absolute force-velocity relationship shows that blebbistatin reduces velocity progressively with increasing concentration. B) Relative force-velocity relationship, normalized to maximal velocity and force, demonstrates a consistent shape across different blebbistatin concentrations, indicating that power generation is similarly affected at all sarcomere lengths.

We performed a series of contractions to derive a force-velocity relation in the myofibrils before and after treatment with blebbistatin. As far as we know, this is the first report for the FV relation in isolated skeletal muscle myofibrils, even in control conditions. As expected, the FV relation showed an inverse hyperbolic relation, and described initially in the classic study by Hill (22). It was also similar to results reported for isolated myosin-actin filaments (6, 46) and entire muscle fibers. We fitted the curves with the Hill equation (see methods) and extracted the Hill coefficients (a/Po). They were comparable to other studies in the literature (6, 36, 46) with values of 0.32 ± 0.07 before treatment (control). After blebbistatin the values decreased to 0.39 ± 0.04 (2 µM blebbistatin) and 0.43 ± 0.06 (5 µM blebbistatin). Blebbistatin affected the FV relation preferentially at low velocities (p = 0.003) and when the curves were normalized, the curvatures appeared similar in all conditions investigated.

## Discussion

In this study, we found that increasing concentrations of blebbistatin (i) decreased the maximum velocity of shortening produced by myofibrils and the myosin-induced actin sliding velocity, (ii) decreased the force produced by myofibrils and isolated myosin filaments, (iii) decreased the curvature of the force-velocity relation in myofibrils. Furthermore, UV radiation reduced the effect of blebbistatin on both velocity and force, but this effect was partially preventable if blebbistatin interacted with myosin prior to exposure to UV light. Altogether, these results show that blebbistatin alters the mechanical properties of myosin from the myofilament to the myofibril levels.

Molecular simulations indicate that blebbistatin binds in the aqueous cavity of the myosin head, between the nucleotide pocket and the actin-binding interface (28), near the P_i_ binding cleft (2). Blebbistatin causes a decrease in the ATP hydrolysis (67), and attaches preferentially in the intermediate state of the myosin-ATPase cycle, in the myosin.ADP.P_i_ position (28). As a result, blebbistatin stabilizes the complex myosin·ADP·P_i_ into a pre-power-stroke state (70). These characteristics of blebbistatin result in significant effects on contractility in skeletal muscles. Blebbistatin decreases the isometric force of permeabilized muscle fibers, as well as stiffness, although to a lower extent than force (41), suggesting that blebbistatin does not prevent myosin attachment to actin, but inhibits the myosin power-stroke. Similar findings have been observed with the myosin inhibitor BDM (8, 23, 41, 42). Although the effects of the drugs are different, they both inhibit P_i_ release from the active site of myosin (74).

Previous studies have shown that the shortening velocity decreases in permeabilized rabbit psoas muscle with 20 μM blebbistatin administration by 45% (65). In the IVMA, we had previously shown a reduced sliding rate proportional to the blebbistatin concentration (49). We corroborate this result and show a proportional decrease in velocity, with the steepest velocity loss between 1 and 5 uM blebbistatin. In our study, we observed that increasing concentrations of blebbistatin led to a proportional decrease in both force and velocity in myofibrils. These observations align with previous reports on the dose-dependent effects of blebbistatin. For instance, a study by Farman et al. demonstrated a dose-dependent reduction in twitch force and sarcomere shortening in isolated rat cardiac myocytes. The study showed that 1 μM blebbistatin induced a slow onset of sarcomere shortening inhibition, while 10 μM blebbistatin resulted in a faster rate of inhibition, highlighting a clear dose-dependency (16). Furthermore, the concentration-dependent reduction in force observed in single rabbit psoas myofibrils are in accordance with our findings, where blebbistatin reduced force and HMM S1 binding kinetics at rates between 0 μM, 7.5 μM, and 50 μM blebbistatin (28). These comparisons highlight the consistency of blebbistatin’s inhibitory effects on myosin function across different systems and concentrations, further validating our results.

We also showed for the first time that the velocity produced by myofibrils was decreased after treatment with blebbistatin. The decrease in myosin-induced actin sliding velocity decrease in IVMA is not as easily explained as the decrease in force. One would assume that blebbistatin could induce a decreased actin detachment rate, (23, 76) although a decreased attachment rate may also be at play (66). In a previous study assessing the relationship between velocity and [MgATP], which is determined by cross-bridge cycle kinetics and the myosin step length (17) Rahman et al. (49) observed a nucleotide concentration-dependent reduction in velocity, where there was an increased effect of blebbistatin on velocity at [MgATP] >0.1 mM and less of an effect at lower concentrations (49). The authors concluded that the blebbistatin induced an accumulation of pre power stroke cross bridges in this state, acting as a break on the actin motility. The rigor or strong attached actomyosin state has a dominating role in limiting velocity at low [MgATP]. At low [MgATP] there is a high likelihood of an increasing number of rigor attached cross-bridges. Accordingly, both with and without blebbistatin there was the same effect on velocity, showing that blebbistatin does not affect the crossbridge at this point in the cycle. While blebbistatin’s effects remain consistent among various experimental setups, questions remain regarding blebbistatin’s effects on regulated and unregulated myofilaments. This discrepancy may be explained by considering the effects of thin filament regulation. Para-amino-blebbistatin (AmBleb) has been shown to reduce the activation of the thin filament by binding to cycling cross-bridges (35). This reduction in thin filament activation might explain why force appears more depressed in myofibrils compared to the myofilament assay. In myofibrils, the presence of regulated thin filaments means that the activation state of the thin filament is crucial for force production, suggesting a degree of cooperativity among the regulatory and force components of the thin and thick filament complex. Consequently, a myosin inhibitor’s effect on reducing thin filament activation would lead to a greater reduction in force in myofibrils, where thin filament regulation is intact, compared to myofilament assays where thin filaments might be unregulated.

Considering the role of cross-bridge intermediate states for the effect of blebbistatin, it is also relevant to discuss the potential influence of the catch state in the myosin filaments from the anterior byssus retractor muscle (ABRM) of *Mytilus edulis*. The catch state is characterized by maintenance of force with very low energy utilization and is regulated by proteins like twitchin. Inhibiting the low to high force transitions, as seen with blebbistatin, promotes catch force, suggesting that force intermediaries can be dissociated under certain conditions, allowing new kinetic states to predominate (4). Specific protein inhibitors are critical for deconstructing the components which contribute to biophysical metrics such as force production. By reducing the primary myosin-driven force with blebbistatin, we can observe the contributions of other regulatory components, as described above regarding thin filament involvement. This understanding underscores the need to consider non-myosin proteins and thin filament regulation in muscle mechanics, providing a more comprehensive view beyond the classical myosin-centric perspective.

Understanding how laboratory manipulations, such as UV exposure before or after HMM incubation, affects blebbistatin function is valuable information for studying actomyosin interaction. The importance of such studies has spurred the development of blebbistatin analogs with specific changes to its interaction with UV light (27, 77). Inactivation experiments using varied UV wavelengths (351-543 nm) and power densities (0 – 10 mW/μm^2^) have shown that blebbistatin is rapidly damaged (56). Because UV promotes the generation of free radicals which in turn damage the myosin or actin, this would thus reduce the velocity in the IVMA. Furthermore, it has been suggested that photo-reactivity of blebbistatin creates intermediates which damage proteins over time. In this current study, time frames during which videos were obtained were too short for protein damage to accumulate and reduce IVMA. Rapid exposure to UV light (180 seconds) was sufficient to diminish blebbistatin effectiveness (Figure 2). Furthermore, videos ranging from 100 – 3000 frames with 0.2s per frame in the IVMA did not show a progressive deterioration of the actin myosin interaction (Supplementary figure 1). Interestingly, increases in velocity over 40 minutes imaging periods may suggest that UV-exposed blebbistatin was preferentially damaged before actin-myosin damage accumulated enough to reduce velocity. Furthermore, we would have expected a reduced velocity over time in controls due to oxidation and conversion of ATP to ADP as assay solution is used.

It has been shown that blebbistain does not affect ATP binding sites and the actin binding sites of myosin (28). Furthermore, S1 bound to actin has low affinity for blebbistatin compared to free S1 (28), suggesting that actin-S1 rigor binding and ATP-induced actin-S1 dissociation is unaffected by blebbistatin. This interpretation suggests that blebbistatin is already in the myosin pocket before the experiments with IVMA were run, causing much of the velocity inhibition. However, it has been noted that blebbistatin effectiveness in myosin can be reduced over time as blebbistatin may be exchanged out of the pocket with the surrounding medium (56). Our results support this interpretation; we noted that IVMA velocity increased over time while blebbistatin was in the motile environment. If blebbistatin leaves the binding pocket, motility would be expected to increase over time as blebbistatin leaves the binding pocket and is less likely to be replaced. We saw this in preparations that were left in the dark. Alternatively, motility with blebbistatin could be expected to increase as it periodically accrues UV damage and re-enters the binding pocket, now with reduced effectiveness and thus greater motility over time. Control experiments without blebbistatin did not exhibit changes in motility over time. Blebbistatin treated with light prior to HMM incubation exhibited no change in motility over time. Blebbistatin may have been damaged both in and out of the myosin pocket, and thus if any replacement of blebbistatin occurred within the pocket, there was no change in the effect on velocity. This seems to imply that damaged blebbistatin has a negligible difference in affinity for myosin despite its reduced effectiveness. Conversely, in IVMA where motility was periodically imaged, there may have only been some UV damage in blebbistatin outside of the pocket over time, which resulted in increasing velocity over time as these unprotected blebbistatin replaced the original HMM-protected blebbistatin (Supplementary Figure 1). The effect of the myosin binding of protecting blebbistatin against degradation is strong, as we observed a marked protection in function in both force (Figure 2) and velocity in the IVMA (Figure 1). These data further support the idea that the effect of blebbistatin is translatable across assemblages of actin-myosin interactions.

<u>Conclusion.</u> Our findings show a reduction in velocity in myosin-actin filaments, a reduction in force in filament interactions, and a reduction in both force and velocity in myofibrils, demonstrating that the effects of blebbistatin are consistent across various levels of actin-myosin interaction. Additionally, the study suggests that blebbistatin damage is preventable by ensuring that it is in the myosin pocket and cannot be damaged by UV light when it interacts with myosin head, which has implications for future experiments using blebbistatin.

## Supplementary figures

**Supplementary figure 1.**
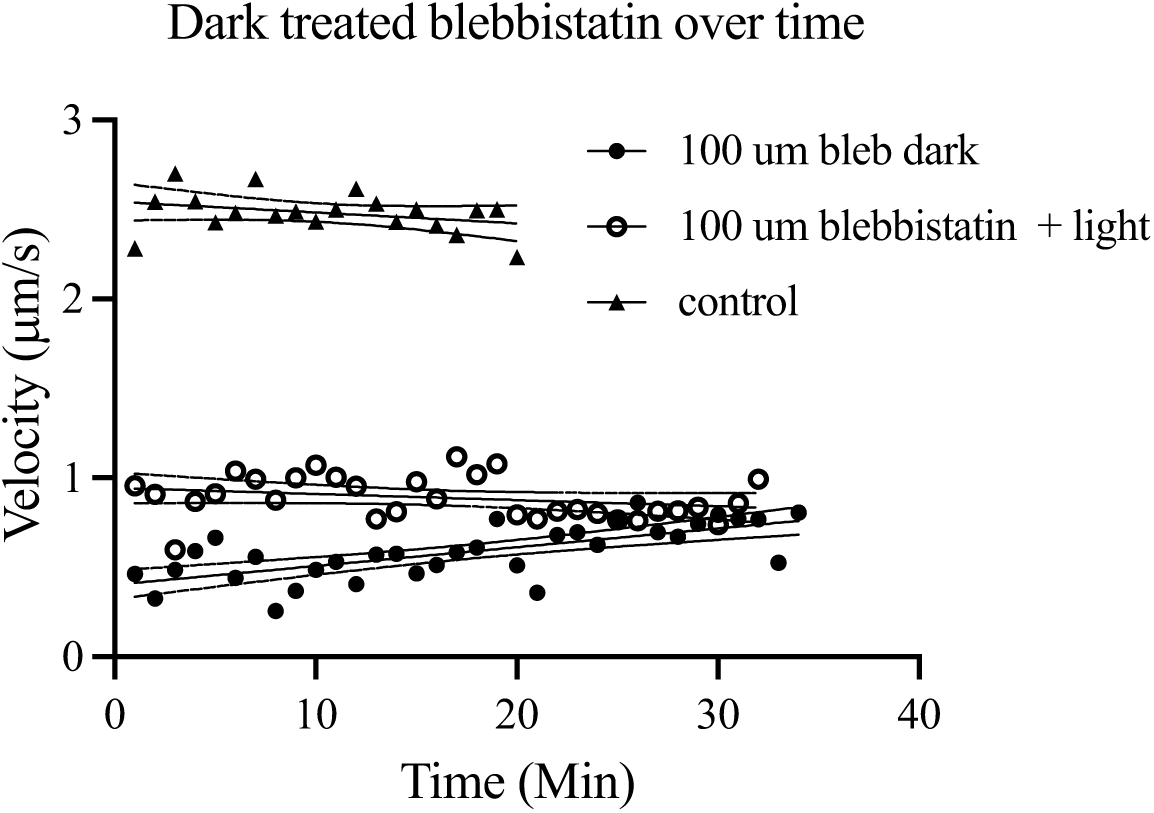
Treatment over time. Control and UV treated blebbistatin maintained efficacy over time. However, blebbistatin left in the dark was less effective over repeated imaging over a period of 40 minutes. This indicates that there may be blebbistatin damage by the UV over time. The blebbistatin may be equilibrating over time and replacing functional blebbistatin, reducing average effectiveness over time. This was not seen when blebbistatin was pre-emptively damaged by laboratory light for 30 minutes prior to 30 minutes of HMM incubation. This shows that blebbistatin was fully damaged by the time motility tracking had begun and could go no lower. A simple linear regression comparing the slopes of the lines revealed that slopes were all significantly different (P < 0.0001), controls and light-exposed blebbistatin slopes were not statistically different from 0. All values shown as mean ± S.E.M.

**Supplementary figure 2.**
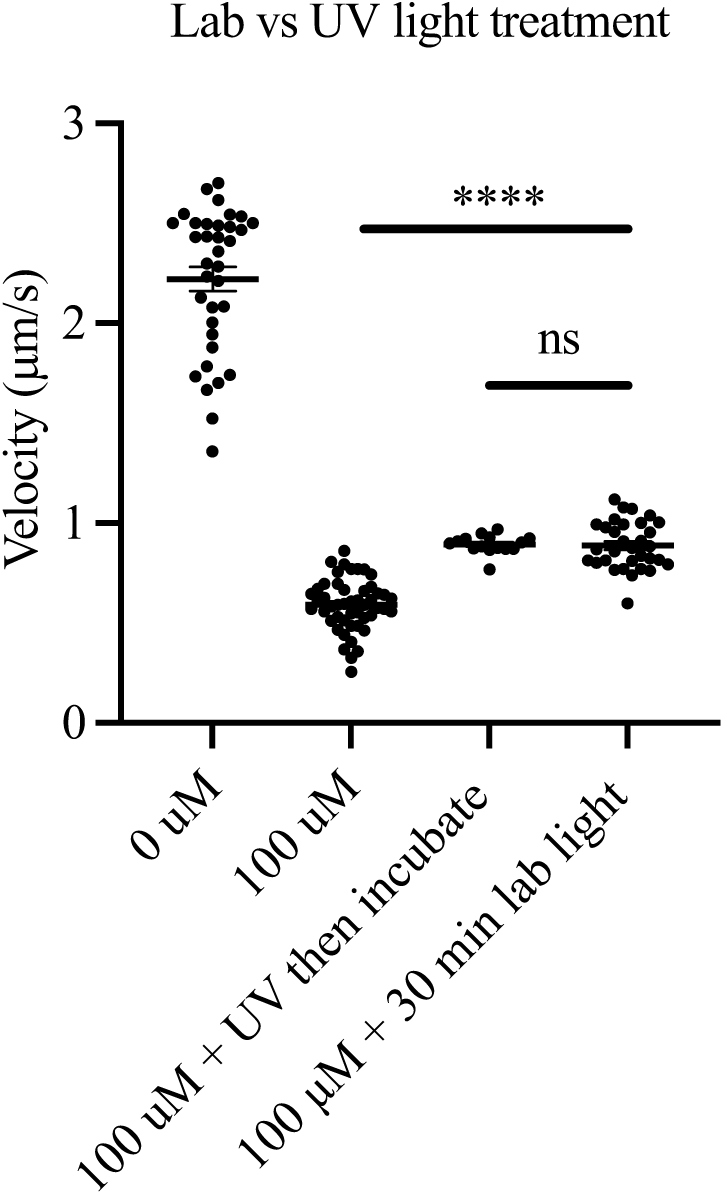
Laboratory light treatment of blebbistatin. **A)** There is a 77% decrease in velocity when 100 *μ*M blebbistatin is added to the HMM relative to controls. When blebbistatin is pre-treated with 30 min laboratory incandescent light, there is a 66% decrease relative to controls. Light treatment caused an increase in velocity relative to no light treatment by 34%. Both light treatments show an increase in velocity relative to 100 *μ*M blebbistatin incubated with HMM in the dark. UV damage was prevented when HMM was incubated before exposure in FFMS as well as in IVMA. The reduction ineffectiveness was proportional in all experiments suggesting that blebbistatin is equally sensitive to blue light in the UV as it is to full spectrum incandescent light. Additionally, this may indicate that diminishment of light-induced damage of blebbistatin reaches a maximal point after which point effectiveness is maintained. All values shown as mean ± S.E.M.

Supplementary Video 1: https://doi.org/10.6084/m9.figshare.22304896.v1 Control FFMS

Supplementary Video 2: https://doi.org/10.6084/m9.figshare.22305898 5 μM FFMS

Supplementary Video 3: https://doi.org/10.6084/m9.figshare.22305946 7.5 μM FFMS

Supplementary Video 4: https://doi.org/10.6084/m9.figshare.22305922 10 μM FFMS

Supplementary Video 5: https://doi.org/10.6084/m9.figshare.22306744 Control IVMA

Supplementary Video 6: https://doi.org/10.6084/m9.figshare.22307359 1 μM IVMA

Supplementary Video 7: https://doi.org/10.6084/m9.figshare.223073715 μM IVMA

Supplementary Video 8: https://doi.org/10.6084/m9.figshare.2444668610 μM IVMA

Supplementary Video 9: https://doi.org/10.6084/m9.figshare.24446695 30 μM IVMA

